# Development of a high-throughput starch digestibility assay reveals wide variation among the A. E. Watkins wheat landrace collection

**DOI:** 10.1101/2022.11.04.515016

**Authors:** Petros Zafeiriou, George M. Savva, Jennifer H. Ahn-Jarvis, Frederick J. Warren, Marianna Pasquariello, Simon Griffiths, David Seung, Brittany A. Hazard

**Affiliations:** Quadram Institute Bioscience, Norwich, UK; John Innes Centre, Norwich, UK

**Keywords:** wheat, landrace, starch digestibility, natural variation, high-throughput, retrograded starch, starch structure, screening tool, pipetting tool, breeding

## Abstract

Breeding for less digestible starch in wheat can improve the health impact of bread and other wheat foods. Based on an established *in vitro* starch digestibility assay by Edwards et al. (2019) we developed a high-throughput assay to measure starch digestibility in hydrothermally processed samples for use in forward genetic approaches. Digestibility of purified starch from maize and wheat was measured using both methods and produced comparable results. Using the high-throughput assay, we estimated starch digestibility of 118 wheat landraces from the core Watkins collection and found wide variation across lines and elite UK varieties, (20% to 40% and 31% to 44% starch digested after 90 minutes respectively). Sieved flour fractions and purified starch for selected lines showed altered starch digestibility profiles compared with wholemeal flour, suggesting that matrix properties of flour rather than intrinsic properties of starch granules conferred the low starch digestibility observed.

## Introduction

Wheat (*Triticum aestivum* L.) is one of the most widely consumed crops worldwide and provides up to 50% of the calories in the human diet, so improving its nutritional quality could deliver health benefits to a large number of people (Shewry and Hey, 2015, Hazard et al., 2020). Reducing the rate and extent of starch digestibility in wheat-based foods can help to maintain healthy blood glucose levels, which is important for the prevention and management of obesity and chronic diseases like type II diabetes (Blaak et al., 2012). Furthermore, there is substantial evidence that consumption of resistant starch, that is the starch that escapes digestion in the small intestine and reaches the colon, can reduce blood glucose levels and help to maintain a healthy gut (Corrado et al., 2022, Belobrajdic et al., 2019, Hughes et al., 2021). Thus, breeding for wheat with less digestible starch is an important strategy to develop healthier staple foods.

A mature wheat grain is predominately starch, 60-70% (w/w), which is the greatest contributor to calories, 10-15% (w/w) protein and 11-15% (w/w) dietary fiber (Shewry et al., 2013, Delcour and Hoseney, 2010). Starch is produced in the endosperm of wheat during grain filling and is composed of two distinct α-glucan polymers, amylose (linear α-1,4-linked chains) and amylopectin (α-1,4-linked chains with α-1,6-linked branches). In their native state, starch polymers exist as partially crystalline granules, which are difficult for humans to digest; thus starch-based foods are typically cooked prior to consumption. Heating starch in the presence of water leads to starch gelatinization making starch polymers more digestible (Wang and Copeland, 2013, Chung et al., 2006, Parada and Aguilera, 2009). Subsequent cooling leads to retrogradation in which starch polymers form crystalline structures that are less digestible (Wang et al., 2015, Corrado et al., 2020). Other intrinsic properties of starch, like its molecular structure can also influence starch digestibility, including the length of its glucan chains and amylose-to-amylopectin ratio (Hazard et al., 2015, Schönhofen et al., 2016, Anugerah et al., 2022).

To date, reverse genetic studies in wheat have demonstrated potential for increasing resistant starch levels using induced mutations in starch biosynthesis genes (Botticella et al., 2018, Schonhofen et al., 2017, Hazard et al., 2015, Fahy et al., 2022). However, initial analyses of some mutants have shown detrimental effects on yield and on pasta and bread-making quality, thus identifying additional sources of genetic variation for starch digestibility could support the development of improved traits for commercial breeding applications (Schonhofen et al., 2017, Hazard et al., 2015). Wheat landraces, locally adapted lines that have not been modified through modern breeding techniques, present reservoirs of genetic diversity that can be introduced into modern cultivars. Of special note is the A.E. Watkins bread wheat landrace collection (Wingen et al., 2014) encompassing 826 bread wheat landraces collected in the 1920s and 1930s from a wide geographic distribution. This collection contains useful variation for a number of agronomic traits and has been used to identify resistance genes for a variety of diseases (Dyck and Jedel, 1989, Bansal et al., 2011, Bansal et al., 2013, Burt et al., 2014). The Watkins lines have been purified by single-seed descent from which many genomic and genetic resources have been developed; a core set of 118 accessions was used to generate nested association mapping populations, all of which were genotyped, have genetic maps available, and are free to access (http://wisplandracepillar.jic.ac.uk/) (Wingen et al., 2017).

Despite the availability of diverse wheat germplasm resources like the Watkins collection, forward screening approaches for starch digestibility have been limited due to the lack of informative, accurate, and efficient phenotyping methods. Screening based on amylose content, which has a positive association with resistant starch content, can identify lines with high levels of resistant starch but cannot identify factors beyond amylose content that may cause resistance to digestion (Chen et al., 2017, Mishra et al., 2016). Only a few studies have developed methods to screen large populations for starch digestibility and they have focused on analyzing purified starch (Wang et al., 2022). However, other components of the wheat flour matrix and processing (e.g. starch retrogradation) could potentially impact starch digestibility (Edwards et al., 2014, Sissons et al., 2021, Qi et al., 2022). To our knowledge no previous studies have screened flour samples of germplasm collections.

Recently, a new *in vitro* method for measuring starch digestibility was published which produces results that are well correlated with the glycaemic response to foods (Edwards et al., 2019) (Figure 1). This method has proven useful for measuring starch digestibility in mechanistic studies (Edwards et al., 2019) and testing early-stage food products. However, modifications were made to tailor the method for screening large wheat germplasm collections accurately and efficiently. Here we describe the development of a high-throughput starch digestibility assay based on the Edwards et al. (2019) protocol, which utilizes a 96-sample format for simultaneous analysis over a 90 min starch digestion. Using standard samples of wheat and maize starch, we validated the assay by comparing starch digestibility profiles to those produced by the Edwards et al. (2019) protocol. The high-throughput assay was then used to screen processed flour samples of the core Watkins (c.Watkins) collection (Wingen et al., 2014) and commercial wheat lines (for bread, biscuit, and animal feed) which revealed natural phenotypic variation for starch digestibility.

**Figure 1.**
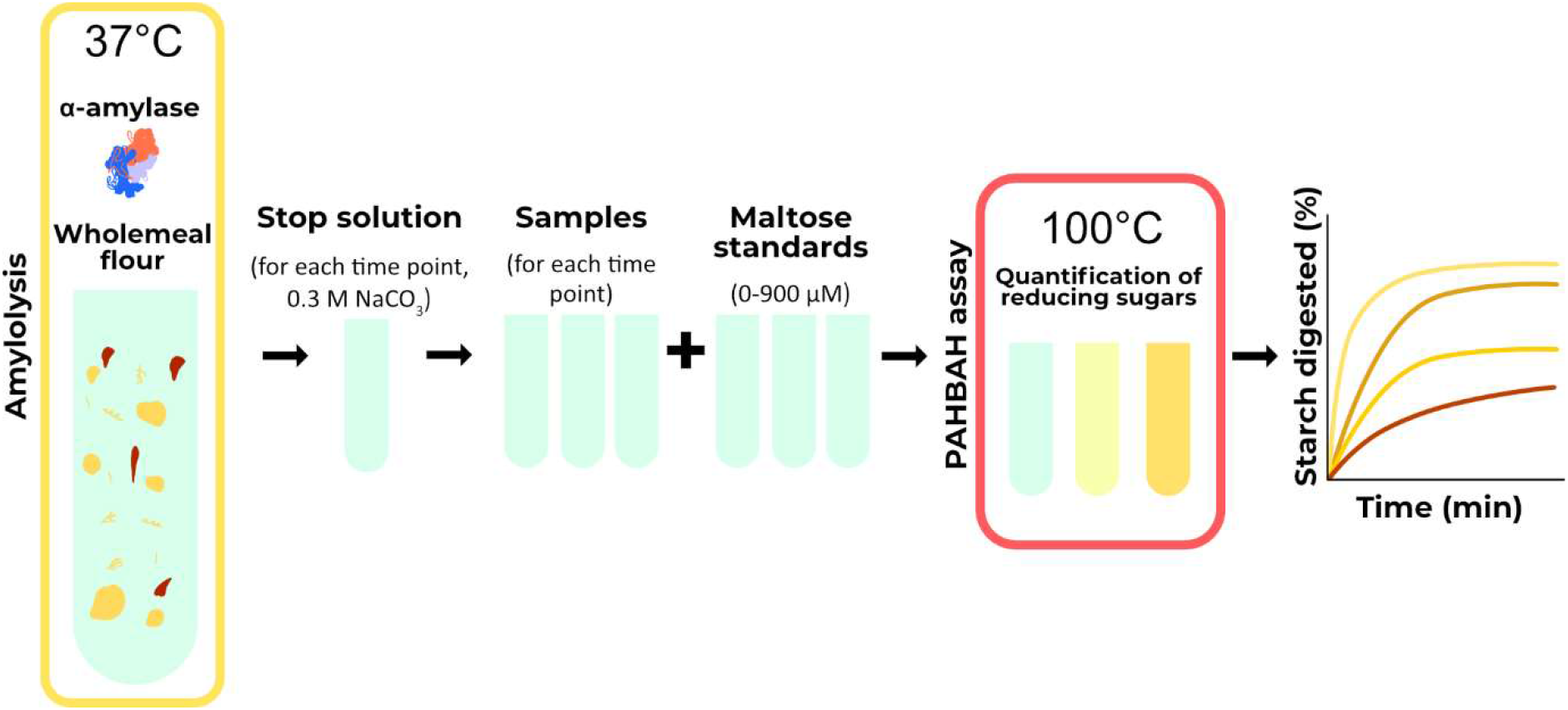
Principle of the in vitro starch digestibility method. For amylolysis, a known enzyme-substrate ratio is used, and starch is hydrolysed by porcine pancreatic α-amylase to produce reducing sugars. During amylolysis, aliquots are transferred to a ‘stop solution’ at predetermined time points to inactivate amylase activity. The reducing sugar concentration is quantified using a colorimetric p-hydroxybenzoic acid hydrazide (PAHBAH) assay and maltose standards (Lever, 1972). The portion of starch digested for each timepoint is calculated based on reducing sugars and is then displayed against time.

## Materials and Methods

### Chemicals

Chemicals used in this study: Percoll™ (17-0891-01, GE Healthcare), 4-Hydroxybenzhydrazide (PAHBAH) (5351-23-5), TRIS (77-86-1), EDTA (60-00-4), SDS (151-21-3), DTT (3483-12-3), Phosphate buffered saline (PBS) (P4417-100), Sodium Carbonate (497-19-8), DMSO (67-68-5), maltose (6363-53-7), sodium hydroxide (1310-73-2) and α-amylase (DFP Treated, Type I-A, saline suspension, 647-015-00-4) were purchased from Sigma-Aldrich Company Ltd., Poole, UK.

### High-throughput starch digestibility assay

Starch digestion assays were carried out on samples that were gelatinized and cooled to accelerate retrogradation following a protocol by Edwards et al. (2019), modified for screening a large number of samples. Wholemeal flour samples were weighed (6 mg) and transferred into a deep well plate (96/1000 μL, Eppendorf). Phosphate buffered saline (600 μL, PBS, pH 7.4) was added to each sample with a 1 mm glass ball to improve mixing. The deep well plate was sealed and secured with a Cap-mat (96-well, 7 mm, Round Plug, Silicone/PTFE) and added to a preheated thermal mixer (80°C) with a 96 SmartBlock™ DWP 1000n attachment (Thermomixer C, Eppendorf Ltd., Stevenage, UK) for 15 min at 1500 rpm to gelatinize the starch, then cooled at 4°C for 21 h to accelerate retrogradation. The plate was then briefly spun (100g for 1 min) to collect condensed liquid on the Cap-mat and placed on the thermal mixer for 30 min at 37°C, 1600 rpm. Time zero samples (50 μL) were collected using a 12-multichannel pipette and transferred into a 2 mL deep well plate containing 1.95 mL of stop solution (17 mM NaCO_3_). Digestion was started by adding pancreatic α-amylase suspended in PBS targeting 2U/mL activity into the samples. Enzyme activity was determined by applying the starch digestibility assay on gelatinized potato starch and obtaining the linear rate of maltose release every 3 minutes (mg/mL). Aliquots (50 μL) were then taken after 6, 12, 18, 24, 40, and 90 minutes from the onset of digestion and transferred to the stop solution.

The stopped reactions were then centrifuged at 1500g for 5 minutes to avoid transferring any starch remnants, and 50 μL of the supernatant was transferred to a new deep well plate; 50 μL of maltose standards (5 - 1000 μM) were also added to the plate. The PAHBAH reducing end assay was used to quantify the reducing ends released (Lever, 1972). Briefly, 0.5 mL of freshly prepared reagent (2 g of p-hydroxybenzoic acid hydrazide dissolved in 38 mL of 0.5 M HCl and 360 mL of 0.5 M NaOH) was added to each sample. The deep-well plate was held at 100°C for 7 min and then placed in an ice bath for 10 min. Samples were then transferred to a microplate, and absorbance was measured at 405 nm using a microplate reader (Bio-Rad Benchmark Plus, Waukegan, Illinois, USA). Reducing sugars were expressed as maltose equivalents, using a standard curve of maltose standards (5 - 1000 μM) from each sample plate. Starch digestibility (%) was expressed according to Edwards et al. (2019); each timepoint’s maltose equivalents were corrected by subtracting the baseline maltose (time zero) and then divided by the maltose equivalent of total starch. Four technical replicates were used, each carried out on a different day.

### Validation of the high-throughput starch digestibility assay

Starch digestion profiles from the high-throughput assay and the Edwards et al. (2019) protocol were compared using standards of purified starch from standard maize, waxy maize, and high-amylose maize (purchased from Merck, formerly Sigma-Aldrich, Darmstadt, Germany), standard wheat starch (purchased from Merck, formerly Sigma-Aldrich, Darmstadt, Germany), and two high-amylose starches: *sbeII* and *ssIIIa*, previously characterized respectively by Corrado et al. (2022), Fahy et al. (2022). Starch samples were aliquoted to make 5.4mg of starch/mL of PBS, and α-amylase activity was adjusted at 2U/mL. The thermal treatment procedure was followed as described above for the high-throughput starch digestibility assay. Three runs were obtained for both protocols over different days. For each run, six replicates per starch sample were placed randomly in the 96-well plate for the high-throughput assay, and two replicates were used for the standard protocol. For each run, starch samples and enzyme solution were prepared in stock and aliquoted for use in both the high-throughput and standard method.

### Field trial design

Grains for the c.Watkins lines were ordered from the Germplasm Resources Unit (John Innes Centre in Norwich, UK) using the publicly accessible SeedStor system https://www.seedstor.ac.uk/; permission to use the materials for research purposes was obtained. The 118 c.Watkins lines were grown in Autumn 2018 in 1 m^2^ plots (one plot per line, except for the low yield lines) at Church Farm, Norfolk UK (52°37’49.2” N 1°10’40.2” E) using standard agronomic practices (Supplemental Figure 1). Based on yield data from previous years, lines with a lower yield performance were grown in duplicate or triplicate (to ensure production of sufficient grains), and grains were pooled for analysis (Supplemental Figure 2). Seeds from elite varieties of bread, biscuit, and animal feed commercial groups of the UK Agriculture and Horticulture Development Board Recommended List (Cougar, Crusoe, Dickens, Diego, Myriad, Paragon, Santiago, Skyfall) were kindly provided by Brendan Fahy (Fahy et al., 2018) (ahdb.org.uk). Elite varieties were grown in 2013 in plots (one per genotype) of 1.5m^2^ at Morley Farm, Norfolk, UK (52°33’15.57” N 1°10’58.956” E).

### Milling and sieving

Grains from the c.Watkins and elite varieties were coarsely milled in a cyclone mill fitted with a 0.5 mm screen (UDY Corporation). Milled samples were passed through a 0.3 mm sieve (Endecotts Limited, London) to produce ‘wholemeal’ flour samples, and selected lines were passed through a 0.053 mm sieve to produce ‘sieved’ flour samples. The flour samples were kept in a vacuum desiccator for five days before analysis.

### Starch isolation

Starch was isolated using an adapted method reported in Hawkins et al. (2021). Wheat flour samples were resuspended with water and filtered through a 100 μm cell strainer (BD Falcon #352360). Samples were then centrifuged at 3000g for 5 min, and the pellets were resuspended in 2 mL of water. The starch suspensions were then overlayed into a Percoll solution (90% v/v) and centrifuged at 2500g for 15 minutes to remove cell walls and proteins. The recovered starch pellets were washed with 1 mL buffer (50 mM Tris-HCl, pH 6.8; 10 mM EDTA; 4% SDS; and 10 mM DTT), transferred into a 2 mL tube and left to incubate for 5 minutes. The starch suspension was then centrifuged at 4000g for 1 min. The pellets were recovered, and the washing procedure was repeated once more. The pellets were then washed three times with 1 mL of water, then once with 100% ethanol. Samples were then kept one day in the fume hood, followed by five days in a desiccator containing silicon dioxide prior to analysis.

### Total starch assay

Wholemeal flour samples were weighed (~8 mg) and transferred into a deep well plate (96/1000 μL, Eppendorf), each well containing 20 μL of DMSO and a 3 mm glass ball to improve mixing. The plate was mixed for 5 min at 1600 rpm to disperse the samples before adding 500 μL of a thermostable α-amylase to each sample. The thermostable α-amylase was solubilized at 1:30 (v/v) in 100 mM sodium acetate buffer, pH 5.0 (Total Starch hexokinase kit, AOAC Method 996.1 1; Megazyme, Bray, IE). The deep well plate was sealed and secured with a Cap-mat (96-well, 7 mm, Round Plug, Silicone/PTFE). Samples were heated at 90°C for 10 minutes at 1600 rpm using a thermal mixer with a 96 SmartBlock™ DWP 1000n attachment (Thermomixer C, Eppendorf Ltd., Stevenage, UK). Total starch content was determined using a Total Starch HK kit (Total Starch hexokinase kit, AOAC Method 996.1 1; Megazyme, Bray, IE) following manufacturer instructions, except volumes of reagents were scaled down by a factor of 10.

### Tools

During optimization of the high-throughput assay, a low-cost 3D-printed pipetting tool was developed, which allowed for manageable weighing and transferring of samples into 96-sample deep well plates and improved speed and control of pipette aspiration (Plate Z). The PLZ 3D design is available to download and print for free (https://www.hackster.io/386082/high-throughput-pipetting-plate-z-bde2c7).

### Data analysis

Statistical analyses and graphs were obtained with R (R version 4.2.1). Datasets of the validation of the high-throughput *in vitro* starch digestibility assay were analysed using the packages lme4 (v 1.1-30) and lmerTest (v 3.1-3) for a mixed model fit (Kunzetsova et al., 2017, Bates et al., 2015, Pinheiro et al., 2022). Plots were made using the ggplot2 package (v 3.3.6) (Wickham, 2016).

For validation of the high-throughput assay, the methods were compared by plotting the estimated starch digestion profiles and by comparing the estimated starch digested at 90 minutes. The bias (difference in estimates between methods) and variation in starch digested at 90 minutes measured by the high-throughput assay was estimated using linear mixed models, including the type of starch as a fixed effect and sample batch as a random effect. The variance was estimated using separate models for each method, while the bias was estimated using a single joint model including all data points, with ‘method’ corresponding to an additional fixed effect.

A linear mixed model including line as a fixed effect and experimental run as a random effect was used for analysis of *in vitro* starch digestibility of the c.Watkins and elite lines, and one-way ANOVA for selected low- and high-digestibility lines as wholemeal flour, sieved flour, purified starch and total starch. The correlation between starch digestibility and total starch was estimated using a linear regression model. All values reported represent the mean, and the number of replicates and variance metrics are specified in the description of the corresponding figure and supplementary datasets. Data sets of starch digestibility and total starch and all statistical analysis code are included as supplementary material.

## Results and Discussion

### Establishment and validation of a high-throughput starch digestibility assay

A key outcome of our study was the development of a high-throughput assay to directly measure starch digestibility of hydrothermally treated flour samples (cooked and cooled wholemeal wheat flour, a processed matrix), based on the Edwards et al. (2019) protocol (referred to as ‘Edwards protocol’ throughout). The high-throughput assay (detailed in the methods section) was scaled from a 6-tube format to a 96-well plate format which required optimizations for maintaining even heat distribution during starch digestions and PAHBAH assays, for the mixing ability of samples, and for recovering adequate sample volumes for analysis. We also developed new tools to allow faster sample handling for sample preparation and pipetting (Plate Z, available free to download from <u>hackster.io</u>). Digestion profiles of maize and wheat starch standards were comparable to profiles generated using the Edwards protocol (Figure 2), particularly with respect to the variance. For example, the difference in the average estimates at 90 mins from the two methods was −0.72 percentage points, and the variation observed at 90 min within the runs was 2.1 percentage points in the high-throughput assay and 2.2 percentage points in the Edwards protocol. There were no significant differences in the percent of starch digested at all the time points measured between the Edwards protocol and the high-throughput assay. Thus, the high-throughput assay provided comparisons between samples that were accurate and reproducible, and the reliability of the assay was deemed sufficient for use as a screening tool to aid in the selection of low- and high-digestibility samples. For future work, it will be important to consider improving the efficiency of upstream steps, including milling, sieving, and weighing samples which present additional bottlenecks.

**Figure 2.**
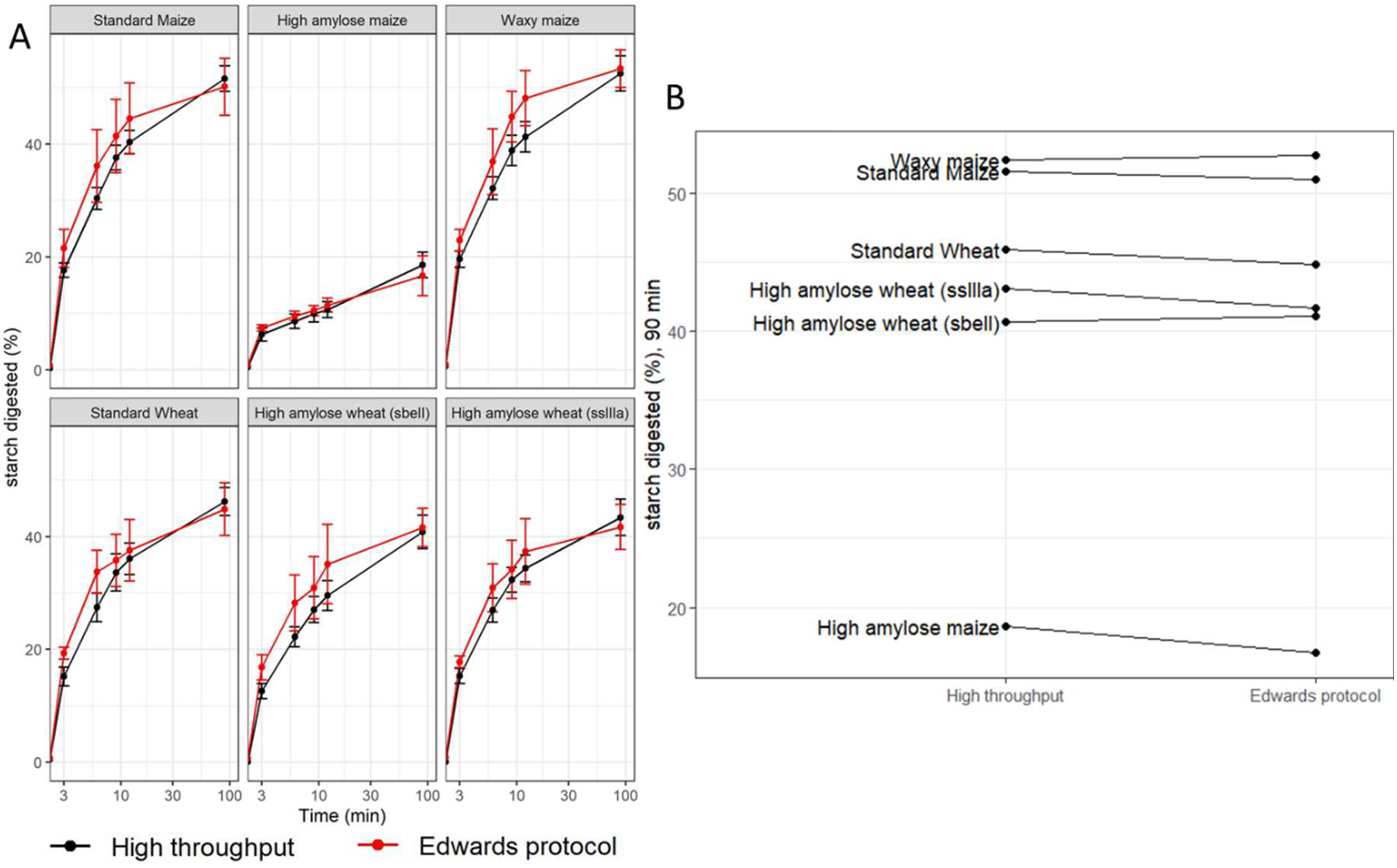
Comparison of the high-throughput starch digestibility assay to the Edwards protocol **A.** digestibility profiles of purified maize and wheat starch produced by the high-throughput assay (black) and the Edwards protocol (red). **B.** Starch digested (%) at 90 min, ranking comparison of the high-throughput assay (left) to the Edwards protocol (right). Values represent the mean ± SD of n = 18 replicates for the high-throughput assay and n = 6 for the Edwards protocol.

Recent studies have reported improved methods to increase the throughput of starch digestibility assays. Wang et al. (2022) developed a 96-sample format assay to screen starch from a wheat MAGIC population and used a multifactorial analysis to identify the most influential factors for starch digestibility (starch granule size distribution, amylopectin chain length distribution, amylose content and endogenous α-amylase activity). Other studies have also aimed to increase the throughput of starch characterization techniques however, the starch properties analyzed may or may not have a direct impact on starch digestibility (Kaufman et al., 2015, Perez-Moral et al., 2018). It is important to consider that other components of wheat flour and factors like processing are likely to have a major impact on starch digestibility thus, determining starch digestibility on processed flour samples may better represent what people may consume in wheat-based foods (Edwards et al., 2014, Edwards et al., 2019, Toutounji et al., 2019, Wang et al., 2022). Some progress has been made for other crops like rice; Toutounji et al. (2019) developed a 15-sample format assay for screening starch digestibility of cooked rice grains with potential to allow analysis of 60 samples per day (if 15 samples are prepared every two hours). In this method, samples were handled individually with a single pipette, which was a limiting factor for the number of samples that could be processed simultaneously. Furthermore, the assay was performed using a set of 8 samples and this assay still needs to be validated on a larger sample set. The high-throughput assay described here addresses some of these key limitations presented by prior assays namely the type of sample analyzed (hydrothermally processed flour) and sample size (96-sample format).

### Screening of c.Watkins landraces and elite varieties

#### *In vitro* starch digestibility

Results of the high-throughput assay revealed a wide range of variation in starch digestibility among c.Watkins landraces; starch digestibility profiles formed a gradient of low to high digestibility rather than two distinct groups, which is expected from complex traits. This variation was evident at all time points (Figure 3). For the c.Watkins lines, the levels of starch digested ranged between 9.7 - 31.6% (at 6 min), 13.2-35% (at 12 min), 14.8-37.2% (at 18 minutes), 16.2 - 36.1% (at 24 min), 19 - 37.8% (at 40 min), and 23.5 - 39.9% (at 90 min). The residual variance from the mixed effect model, measuring the variability between technical replicates from the same sample in the same run, was 3.7 percentage points. The variance using wholemeal flour was higher than the variance using starch standards in the validation experiment, and this could be due to the higher variability in wholemeal flour composition compared to purified starch samples. The number of technical replicates needed to estimate starch digestibility to a given precision can be calculated from residual variance.

**Figure 3.**
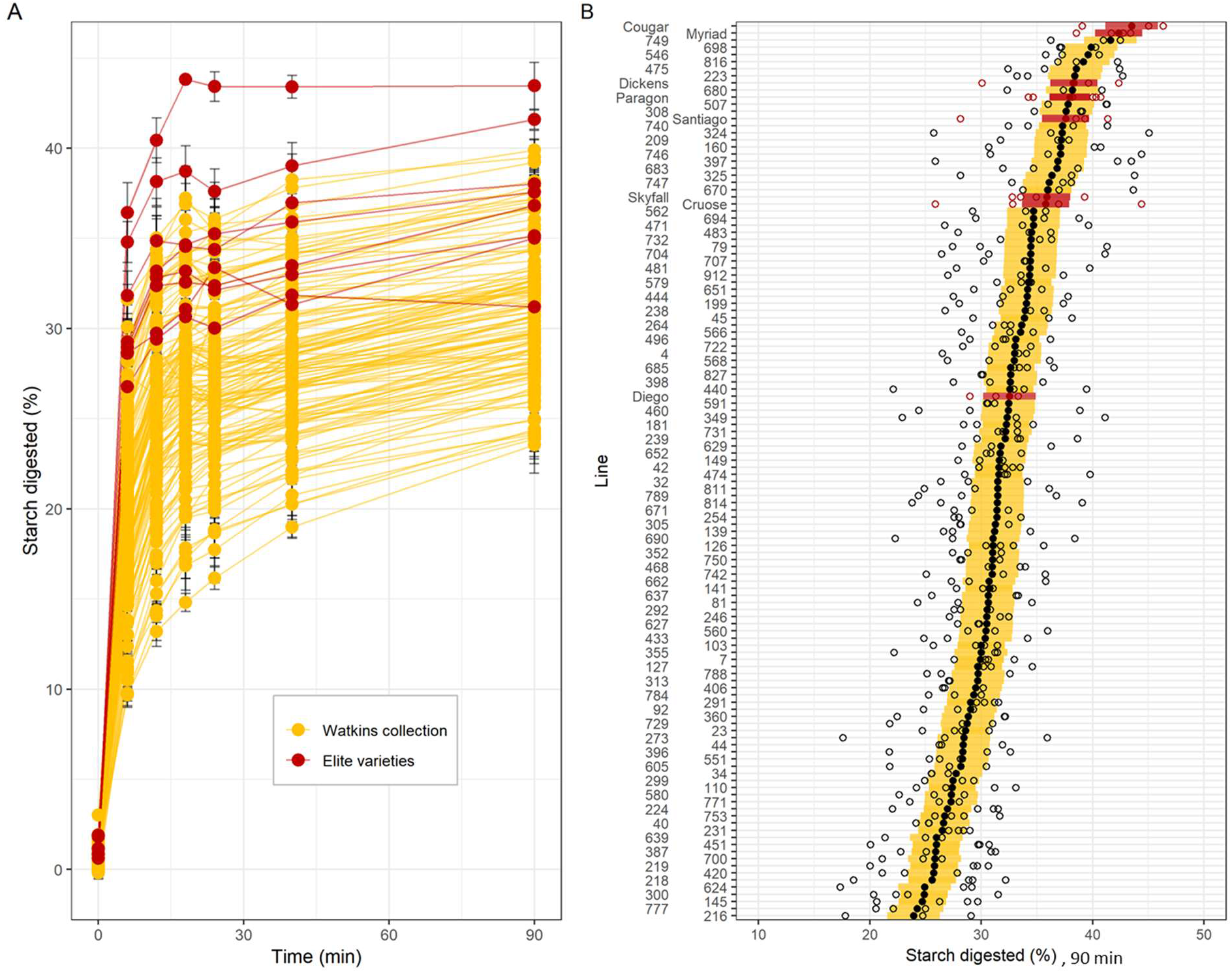
Starch digestibility and total starch of c.Watkins landraces (yellow) and elite varieties (red) of wholemeal flour. **A.** Starch digested (%) for c.Watkins (yellow) and elite varieties (red). Points and lines represent the mean and standard error from between 3 and 6 technical replicates per line. **B.** Starch digested (%) at 90 min. Individual data points from technical replicates are shown in white dots, and marginal mean values from a mixed effects model with line as a fixed effect and experimental run as a random effect are shown in black (c. Watkins) and red dots (elite). The standard error estimated from the model is displayed as yellow (c. Watkins) and red (elite) bars. Values represent mean ± SE of n ≥ 3 replicates.

There was no evidence for a relationship between the plot location in the field and the starch digestibility at 90 min, as no trend was observed between field rows and columns (Supplemental Figure 3). Elite UK varieties showed less variation and greater starch digestibility compared to most c.Watkins lines, although a direct comparison is confounded by the different growing conditions between the two groups. For example, the levels of starch digested for the elite varieties ranged between 26.8 – 36.4% (at 6 min), 29.4-40.4% (at 12 min), 30.6 – 43.8% (at 18 min), 30 – 43.4% (at 24 min), 31.3 – 43.4% (at 40 min) and 31.2 – 43.5% (at 90 min).

Our results demonstrate that natural phenotypic variation for starch digestibility exists in the c.Watkins collection (representing the genetic diversity of the entire Watkins collection, 826 lines) as well as in elite varieties representing commercial groups of bread, biscuit and animal feed groups, thus there is potential for identifying underlying genetic diversity. The levels of variation observed in the c.Watkins were greater than the elite varieties screened, which is consistent with the higher levels of genetic diversity previously reported for the c.Watkins collection compared to modern European bread wheat varieties, as well as increased phenotypic variation for agronomically desirable traits, such as stripe, leaf, and stem rust resistance, as well as grain surface area and grain width (Wingen et al., 2014, Bansal et al., 2011, Bansal et al., 2013, Randhawa et al., 2015, Toor et al., 2013). It is important to note that a limitation of our study was that the grain samples of c.Watkins lines and elite varieties were not produced in the same field trial, and each used a plot per line, so it is possible that environmental conditions between and within fields could influence the average starch digestibility levels and the variation observed; this will be important to consider in future trials. Furthermore, with the availability of structured germplasm panels (nested association mapping populations) and genotype data (Wingen et al., 2017), there is now potential for use of the high-throughput assay to investigate the genetic factors underlying the variation in starch digestibility observed through approaches like QTL analysis which will aid in identification of underlying candidate genes and development of tools for marker assisted-selection.

#### Total Starch

Total starch (TS) content of wholemeal flour varied significantly for c.Watkins landraces (43 ± 3.3 g/100 flour to 61 ± 2.4 g/100 flour, mean ± SE, *p* ≤ 0.001), and the elite varieties (46 ± 3.6 g/100 flour to 61 ± 0.9 g/100 flour, mean ± SE, *p* ≤ 0.05) (Supplemental data). The elite variety Diego had the highest TS content (61 ± 0.9 g/100 flour, mean ± SE), and the c.Watkins line 651 had the lowest (43 ± 3.3 g/100 flour, mean ± SE). Most of the samples had a TS content between 47 to 57 g/100 flour. Linear regression analysis showed that total starch content only weakly correlated (R^2^ = 0.0108) with starch digested at 90 min which suggests that the differences in total starch content cannot explain the variation observed for starch digestibility (Supplemental Figure 4).

#### Analysis of low- and high-digestibility lines as wholemeal flour, sieved flour, and purified starch

Three low-digestibility lines: 777 (WATDE0111), 216 (WATDE0025), and 639 (WATDE0083); and five high-digestibility lines: including two c.Watkins landraces 816 (WATDE0117), 308 (WATDE0042) and the three UK elite varieties Myriad, Dickens, and Paragon were selected for further analysis. The selection was based on the high-throughput starch digestibility assay results, specifically the ranking (low to high), to represent the variation observed in the screen.

Starch digestibility profiles differed considerably for wholemeal flour, sieved flour and purified starch, suggesting that other flour components besides starch likely contributed to the variation in starch digestibility observed (Figure 4). For wholemeal flour (<0.3 mm), starch digestibility profiles of selected ‘low-’ and ‘high-digestibility’ lines grouped separately where the high-digestibility samples varied between 37.6 - 41.6% and low-digestibility samples varied between 23.5 - 24.3% (at 90 min) (Figure 4A). For sieved flour (< 0.05 mm), the digestibility of two low-lines (216 and 639) increased significantly (p < 0.05) compared to wholemeal samples, and the third low-line (777) remained low (Figure 4B). In sieved flour, the high-digestibility samples varied between 38.1 - 42.7%, whereas low-digestibility samples varied between 26 - 35.4% (at 90 min). Purified starch samples also showed a greater extent of starch digested at 90 minutes compared to wholemeal and sieved flour fractions (p < 0.001) (Figure 4B) but there were no differences between the high- and low-digestibility groups in purified starch samples. For example, high-digestibility lines differed between 46.9 - 52.3% and low-digestibility lines varied between 47.6 - 49.2% (at 90 min); the variation observed at 90 min within the runs was 3 SD (95% CI= 2.3 to 4.2%).

**Figure 4.**
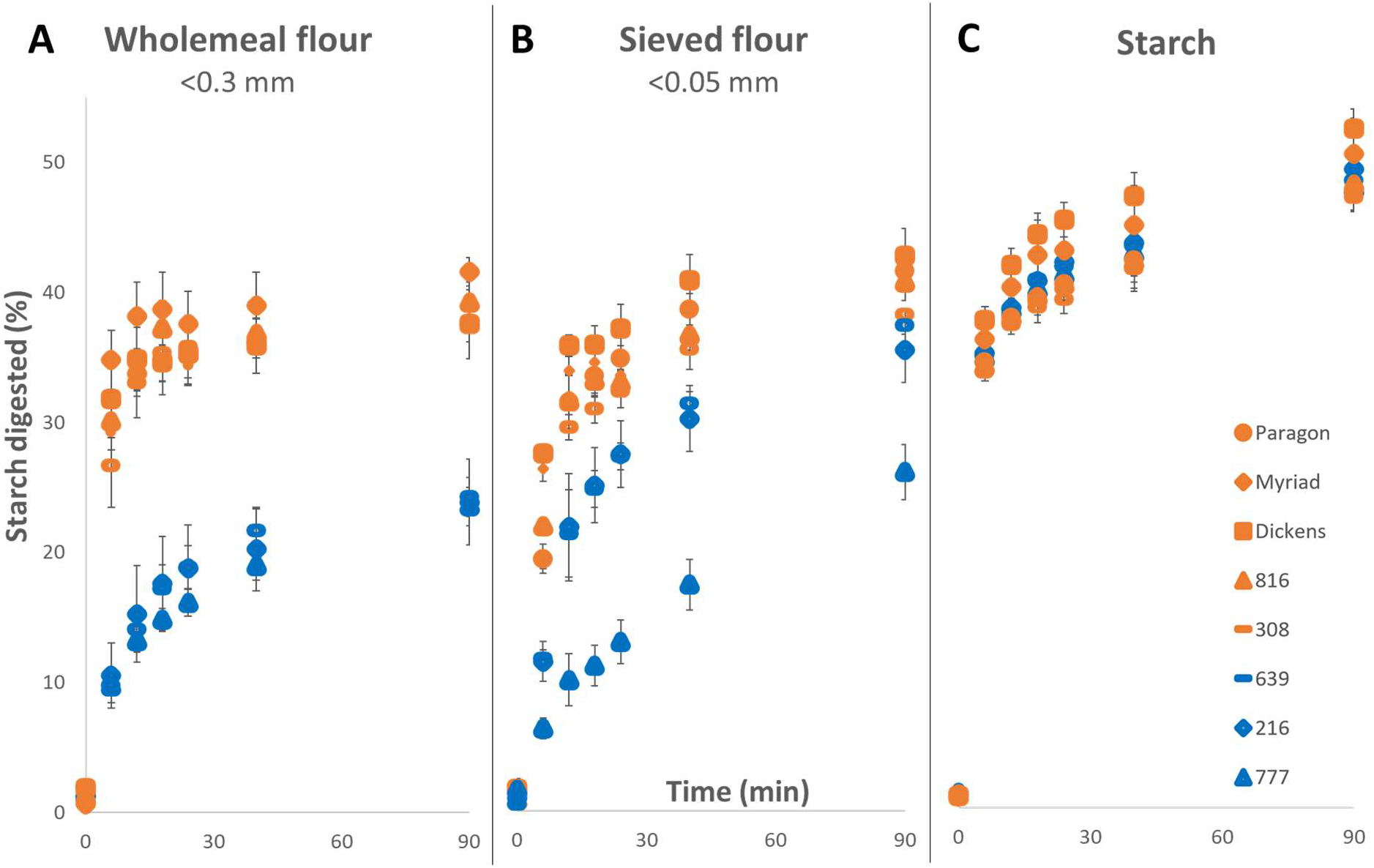
Starch digestibility of selected low- and high-digestibility lines measured in wholemeal flour (A), sieved flour (B) and purified starch (C) using the high-throughput assay. Samples in blue represent the low-digestibility lines, and samples in orange represent the high-digestibility lines. Values represent mean ± SE of n = 3 replicates.

The varying digestibility profiles observed across purification steps suggest that multiple mechanisms in the selected lines could be affecting starch digestibility and that different factors in each line could have a distinct effect. This is consistent with prior studies which have shown effects of flour characteristics and starch structure on starch digestibility in cooked wheat samples including particle size of wholemeal flour, protein content, and chain-length distribution of starch polymers (Obadi et al., 2020, Lin et al., 2020, Sissons et al., 2021). In our study, selected lines were also analyzed for flour properties (flour particle size, endogenous α-amylase and protein content) and starch structural properties (starch granule size distribution and chain-length distribution), however, none of the factors strongly correlated with starch digestibility, which could be due to the high phenotypic variation of the Watkins collection (Supplemental Table 1 and Supplemental Figures 5,6,7). An important limitation of this aspect of the study was the small number of lines selected for this analysis; prior work by Wang et al. (2022) identified starch properties influencing starch digestibility in a screen of 224 wheat starches. Nevertheless, our study suggests that a combination and interaction of many factors are required to achieve low starch digestibility profiles in flour, so using a high-throughput starch digestibility assay for screening or selection approaches may be more efficient and informative than selection based solely on underlying factors such as starch molecular structure.

## Abbreviations

c.Watkins: core Watkins
PAHBAH: p-hydroxybenzoic acid hydrazide
TS: Total: starch

## Authors contributions

Petros Zafeiriou: Conceptualization, investigation, methodology, project administration, data curation, and formal analysis, writing – original draft preparation, writing – review & editing, George M. Savva: methodology, data curation, visualization, formal analysis, writing – review & editing, Jennifer Ahn-Jarvis: investigation, methodology, writing – review & editing, Frederick J. Warren: conceptualization, methodology, supervision, writing – review & editing, Marianna Pasquariello: investigation, methodology, writing – review & editing, Simon Griffiths: conceptualization, methodology, supervision, writing – review & editing, David Seung: conceptualization, investigation, methodology, supervision, writing – review & editing, Brittany A. Hazard: conceptualization, funding acquisition, methodology, supervision, writing – review & editing.

The authors declare no conflicts of interest.

## Acknowledgements

This project was funded by the UKRI Biotechnology and Biological Sciences Research Council Norwich Research Park Biosciences Doctoral Training Partnership [grant number BB/M011216/1] (PZ) and the UKRI Biotechnology and Biological Sciences Research Council (BBSRC) Institute Strategic Programme’ Food Innovation and Health’ (grant number BB/R012512/1) and its constituent projects BBS/E/F/000PR10343 (Theme 1, Food Innovation) and BBS/E/F/000PR10345 (Theme 2, Digestion in the Upper GI Tract) and the BBSRC Core Capability Grant BB/CCG1860/1 awarded to Quadram Institute Bioscience; the BBSRC Institute Strategic Programme Grants’ Molecules from Nature’ — Crop Quality BBS/E/J/000PR9799 awarded to the John Innes Centre and ‘Designing Future Wheat’ BB/P016855/1 awarded to the John Innes Centre.

## Notes

### Competing Interest Statement

The authors have declared no competing interest.

### Summary of Updates

The introduction has been reshaped. Manufacturer details were added for the sieving, and processed starch has been named as gelatinized and cooled instead of gelatinised and retrograded

