## Supplemental material for "Development of a high-throughput starch digestibility assay reveals wide variation among the A. E. Watkins wheat landrace collection" (from another version of this preprint: version 1, 2022-11-04)

| | | |
|---|---|---|
| | 698 | 349 |
| | 694 | 324 |
| | 694 | 313 |
| | 690 | Soison |
| | 690 | 313 |
| | 685 | 308 |
| | Soison | 305 |
| | 685 | 300 |
| | 683 | 299 |
| | 680 | 292 |
| | 671 | 291 |
| | 670 | 352 |
| | 662 | 273 |
| | 652 | 264 |
| | 639 | 254 |
| | 637 | 325 |
| | 629 | 246 |
| | 627 | 239 |
| | 627 | 238 |
| | Soison | Soison |
| 912 | 624 | 231 |
| 827 | 605 | 224 |
| 827 | 591 | Soison |
| 827 | 591 | 223 |
| 816 | 580 | 219 |
| 816 | Soison | 218 |
| 814 | 579 | 216 |
| 811 | 568 | Soison |
| Soison | 568 | 209 |
| 811 | 566 | 209 |
| 789 | 562 | Soison |
| 788 | 560 | 199 |
| 784 | 551 | 199 |
| 777 | 546 | 181 |
| 771 | 507 | 160 |
| 753 | 496 | 149 |
| 750 | 483 | 145 |
| 749 | 481 | 141 |
| 747 | 475 | 139 |
| Soison | Soison | Soison |
| 746 | 474 | 127 |
| 742 | 471 | Soison |
| 742 | 471 | 126 |
| 740 | 468 | 110 |
| 651 | Soison | 103 |
| 732 | 468 | 92 |
| 731 | 460 | 81 |
| Soison | 451 | 79 |
| 731 | 444 | Soison |
| 729 | 440 | 45 |
| 729 | 433 | 44 |
| 722 | 420 | 42 |
| 707 | 406 | 40 |
| 705 | 398 | 34 |
| 705 | 397 | 34 |
| 704 | 396 | 32 |
| 700 | 387 | 23 |
| 700 | 360 | 7 |
| 698 | 355 | 4 |
| Soison | Soison | Soison |

***Supplemental Figure 1.*** *Field trial (1 $m^2$ plots at Church Farm, Norfolk UK (52°37'49.2"N 1°10'40.2"E)) of c.Watkins. Numbers represent the lines, and Soison was used as a standard for the field trial validation.*

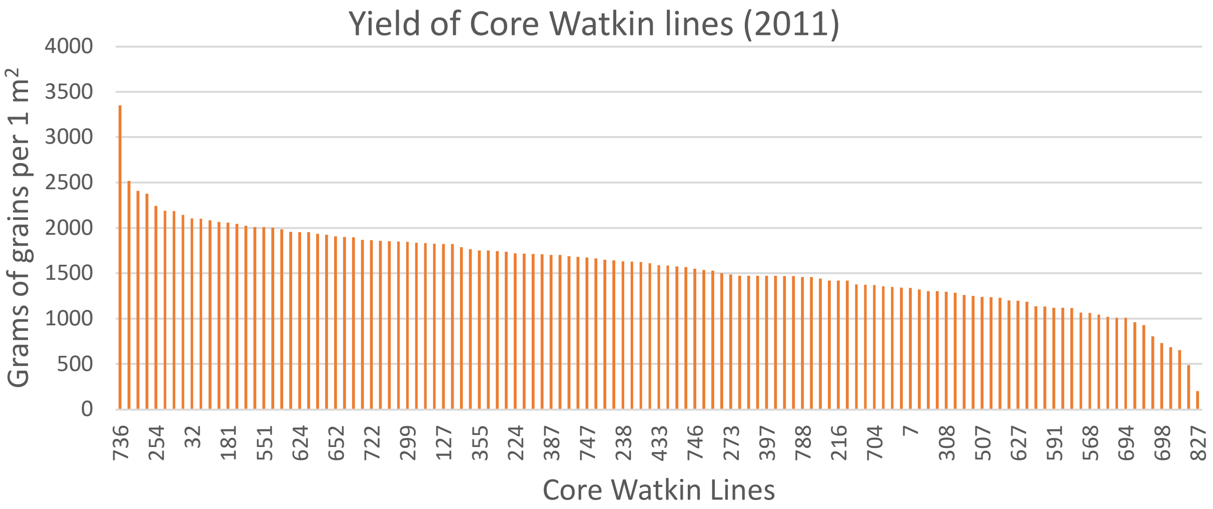


***Supplemental Figure 2.*** *Core Watkins' collection yield (2011). Provided by Germplasm resources unit.*

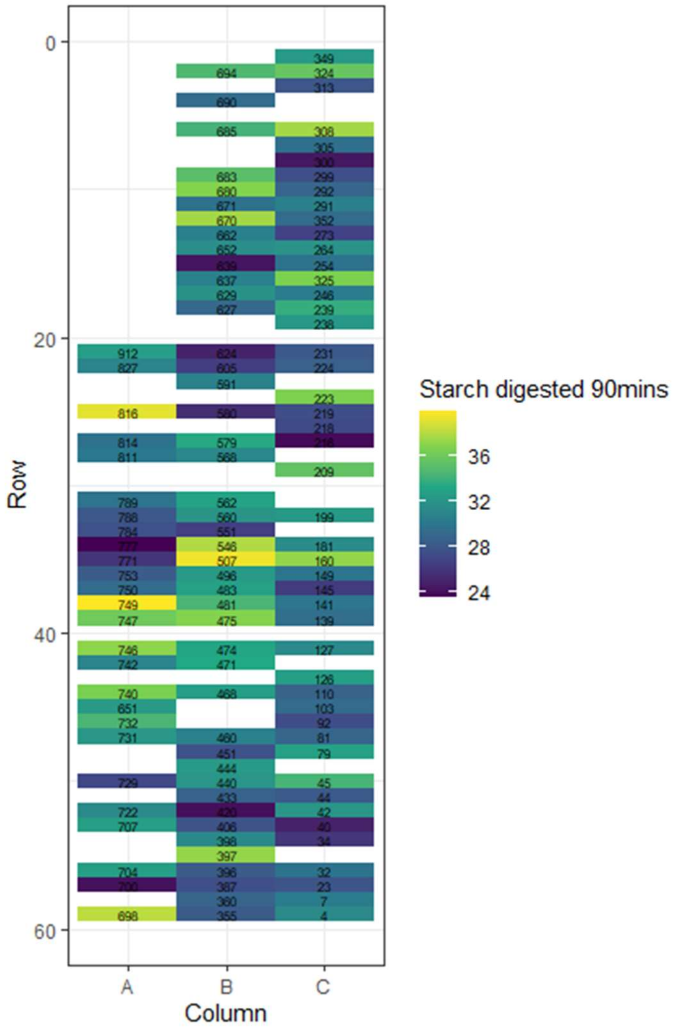


***Supplemental Figure 3.*** *Field design of c.Watkins collection and starch digested (%) at 90 min.*

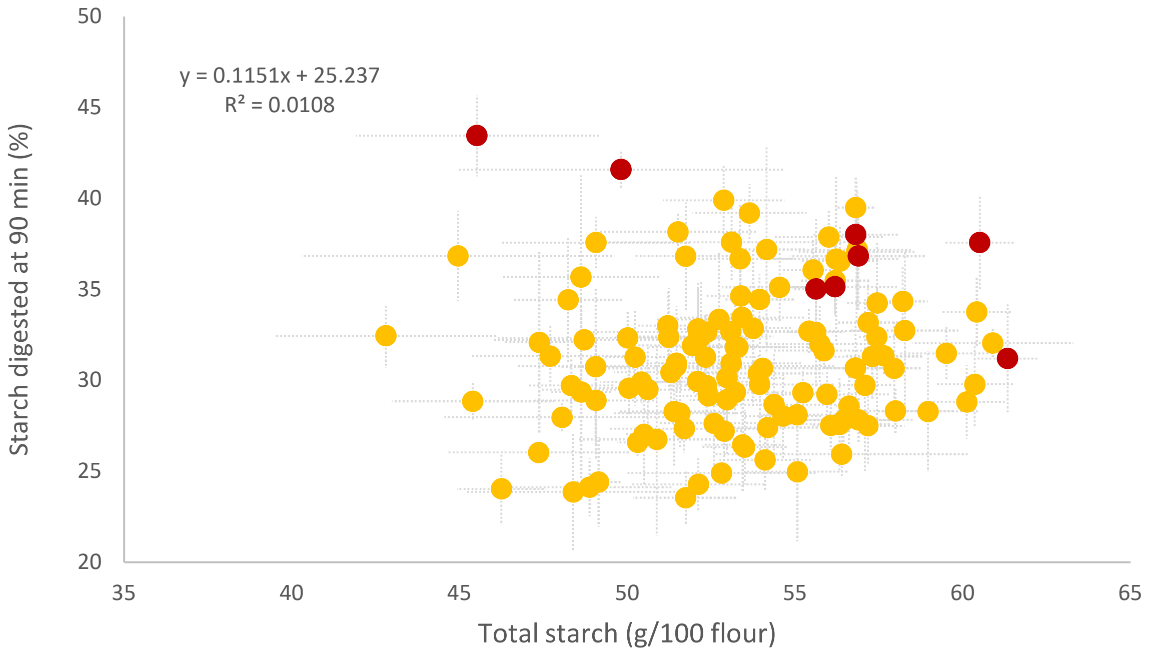


***Supplemental Figure 4***. *Scatter plot of starch digestibility (%) and total starch content (g/100 flour) of c.Watkins landraces (yellow) and elite varieties (red). Error bars represent ± (SE).*

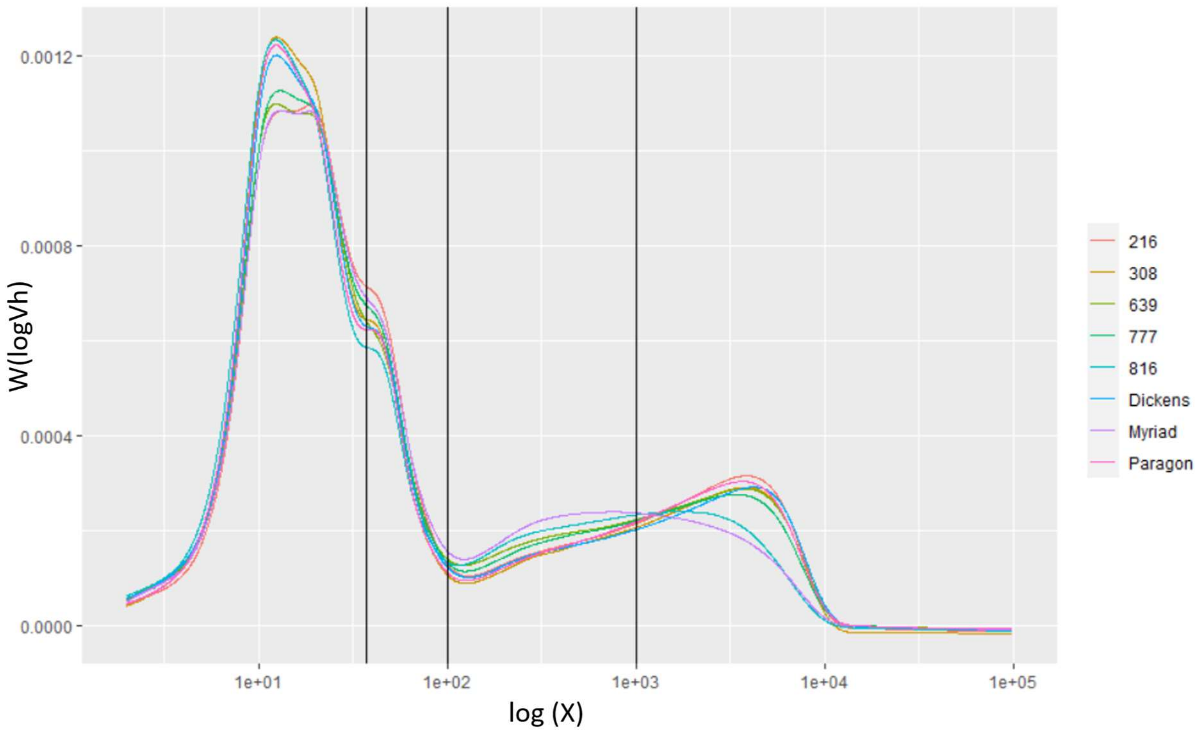


***Supplemental Figure 5.*** *Size exclusion chromatography data on selected wheat lines (216, 308, 639, 777, 816, Dickens, Myriad and Paragon). The Y-axis is expressed as wLogVh; the X-axis is the degree of polymerization and is expressed as log(X).*

***Table 1.*** *Protein content of wholemeal flour was measured on a fresh weight basis. Coulter counter analysis for granule size distribution described as the relative volume of B-type granules (%), B-type granule diameter (µm) and A-type granule diameter (µm). Chain-length distribution was determined by SEC: amylopectin (AP) long to short chains (ratio of 37<x<100 to <37 DP), AM (amylose) long/short chains (ratio of and >1000 to >100-1000 DP), AM/AP (ratio of >100 to <100). Endogenous α-amylase was determined as Ceralpha Units/g of flour. Values represent mean ± SE. Statistically significant value p shows the difference between low digestibility and high digestibility group.*

| | Lines | Protein content (g/100 flour) | B granule (%) | B-type granule diameter (µm) | A-type granule diameter (µm) | AP long/short chains | AM long/short chains | AM/AP ratio | Endogenous α-amylase Ceralpha Units/g flour |
|---|---|---|---|---|---|---|---|---|---|
| p-value | | p < 0.001 | ns | p < 0.01 | p < 0.001 | p < 0.001 | p < 0.005 | ns | p < 0.05 |
| Low Digestibility | 639 | 15.1 (0.17) | 37.1 (1.9) | 6.4 (0.2) | 18.2 (0.1) | 0.23 (0.01) | 0.94 (0.06) | 0.32 (0.01) | 0.06 (0.01) |
| | 777 | 14.2 (0.16) | 34 (2) | 6 (0.1) | 18.2 (0.2) | 0.24 (0.01) | 0.93 (0.05) | 0.30 | 0.07 (0.01) |
| | 216 | 18.4 (0.05) | 28.6 (1.6) | 8.4 (0.2) | 20.1 (0.2) | 0.26 | 1.10 (0.04) | 0.29 (0.01) | 0.15 (0.01) |
| High Digestibility | 308 | 14.9 (0.18) | 35.3 (1.7) | 6.2 (0.1) | 17.7 | 0.21 (0.01) | 1.06 (0.06) | 0.26 (0.02) | 0.13 (0.01) |
| | Myriad | 11.1 (0.22) | 34.2 (2.4) | 6.7 (0.1) | 19.2 (0.5) | 0.25 | 0.84 (0.13) | 0.29 (0.02) | 0.06 (0.01) |
| | 816 | 17.4 (0.19) | 34.8 (0.7) | 6.9 (0.4) | 19 (0.5) | 0.20 (0.01) | 0.65 (0.05) | 0.27 (0.03) | 0.09 (0.01) |
| | Paragon | 13.5 (0.24) | 30.9 (1) | 6.6 (0.1) | 20.1 (0.2) | 0.21 | 1.16 (0.14) | 0.29 (0.01) | 0.05 (0.01) |
| | Dickens | 10.6 (0.15) | 29 (3.3) | 6.3 (0.2) | 19.2 (0.3) | 0.22 | 1.09 (0.04) | 0.28 | 0.05 (0.01) |

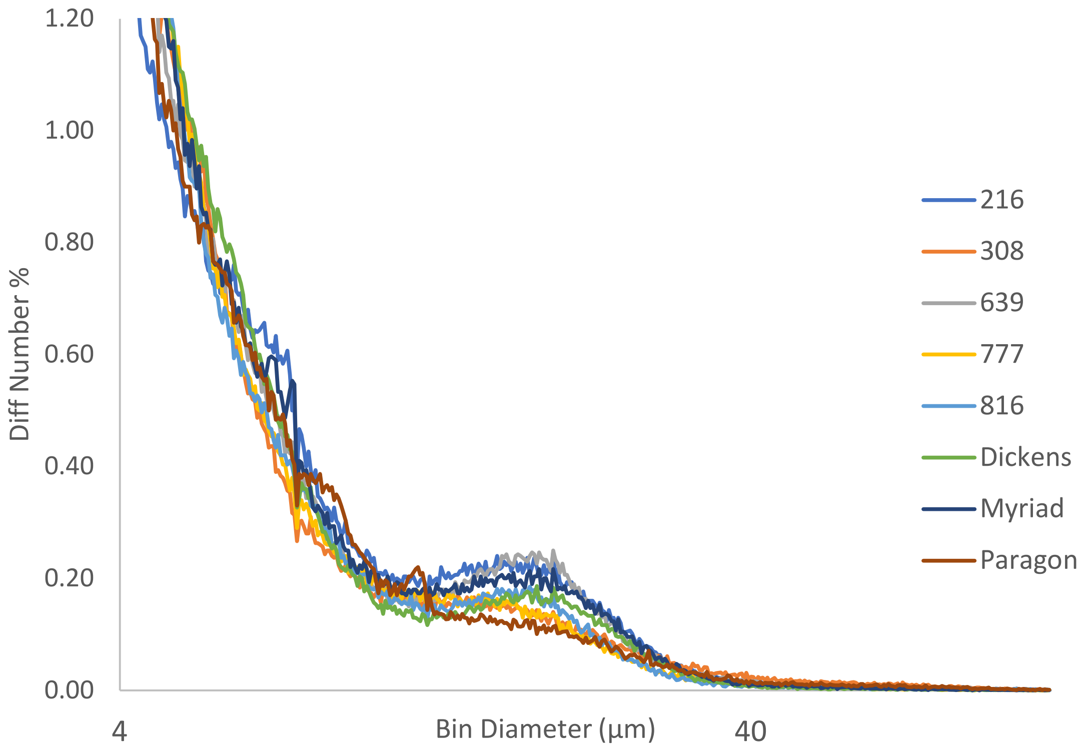


**Supplemental Figure 6.** Differential Particle Size Distribution of wholemeal flour particles by Coulter counter analysis, data is shown on a log scale in the x-axis.

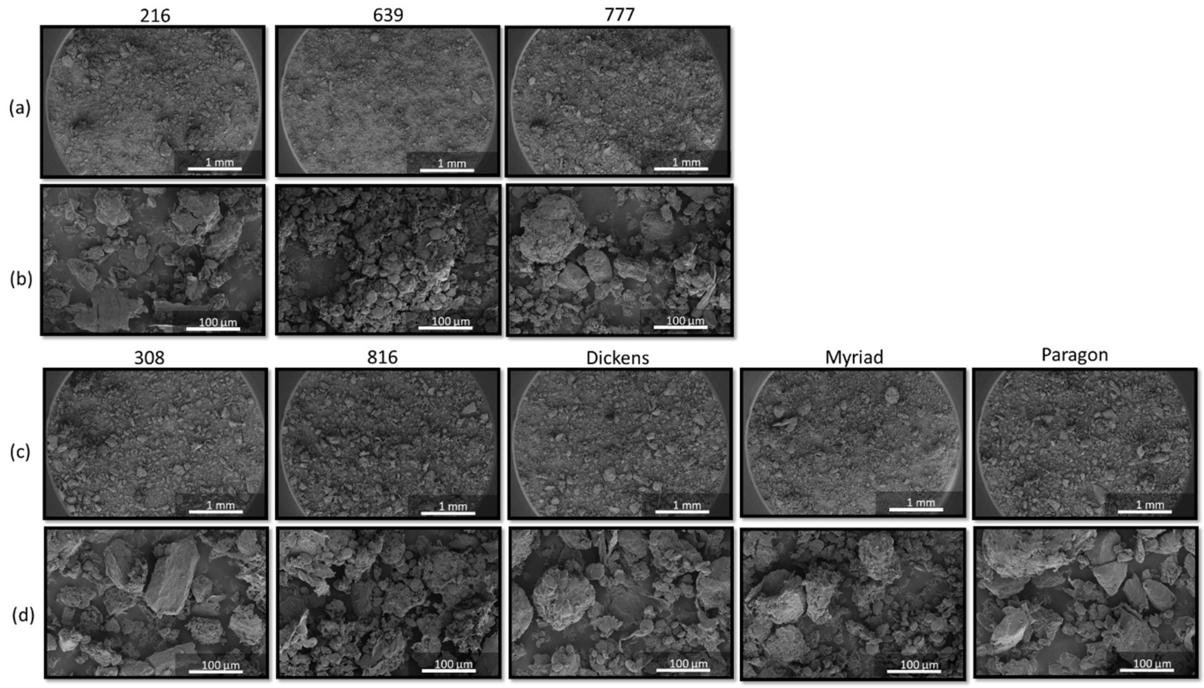


***Supplemental Figure 7.*** *SEM micrographs of wholemeal flour for low- and high-digestibility lines.* ***(a)*** *and* ***(b)*** *are from of low digestibility lines and* ***(c)*** *and* ***(d)*** *are from high digestibility lines.*
